# Homologous recombination delayed repair in oocytes in the bdelloid rotifer *Adineta vaga* post radiation

**DOI:** 10.64898/2026.04.30.722046

**Authors:** Victoria C. Moris, Alexandre Philippart, Cécile Husson, Bernard Hallet, Boris Hespeels, Karine Van Doninck

**Affiliations:** Université Libre de Bruxelles (ULB), research unit in Molecular Biology and Evolution (MBE), Brussels, 1050, Belgium; University of Namur, Research Unit in Environmental and Evolutionary Biology (URBE), Institute of Life, Earth & Environment (ILEE), Namur, 5000, Belgium; Université Libre de Bruxelles (ULB), Brussels Laboratory of the Universe (BLU-ULB), Brussels, 1050, Belgium; Université Libre de Bruxelles (ULB), Marine Biology Laboratory (BIOMAR), Brussels, 1050, Belgium; Université Catholique de Louvain (UCLouvain), The Louvain Institute of Biomolecular Science and Technology, Louvain-la-Neuve, 1348, Belgium

**Keywords:** In situ hybridization, bdelloid rotifer, DNA repair, Homologous Recombination, Non-Homologous End-Joining, radiation, oocyte, germline, somatic

## Abstract

Bdelloid rotifers are known to survive desiccation and high doses of ionizing radiation. This extreme resistance is notably due to their capacity to cope with numerous DNA double-strand breaks (DSBs). Genes encoding key components of the non-homologous end joining (NHEJ) DNA repair pathway are strongly upregulated in the bdelloid rotifer *Adineta vaga* following exposure to ionizing radiation. Considering the notably high doses tolerated by these organisms, their capacity to efficiently restore genome integrity is particularly striking. Although NHEJ is generally regarded as less accurate than homologous recombination (HR), the absence of major genomic rearrangements in the descendants of irradiated rotifers suggests that DNA repair occurs with high fidelity. Terwagne et al. recently reported a delayed repair in germline nuclei, occurring during oocyte development when homologous chromosomes pair, thereby enabling template-based repair through HR. In this study, we established an *in situ* hybridization approach on *A. vaga* cryosections to investigate the spatial and temporal expression of key actors involved in NHEJ, HR, and Base excision repair (BER) pathways in somatic and germline tissues. We show that NHEJ (KU80) and BER-related genes (PARPs) as well as *A. vaga* Ligase E (putatively involved in DNA repair) are expressed early after radiation exposure in the somatic syncytium. In contrast, HR-related genes (Rad51: two paralogs, Rad54), as well as PCNA (involved in DNA replication, NER, BER, HR) are expressed later in maturing oocytes, indicating the activation of a delayed homologous recombination repair pathway in germline nuclei. Nurse cells, which express genes associated with both HR and NHEJ pathways, may rely on both mechanisms for their own DNA repair while also supplying mRNAs to the maturing oocyte. Our results provide new evidence for a differential regulation of DNA DSB repair pathways between soma and germline in bdelloids, with NHEJ predominating in somatic tissues and HR in the germline of *A. vaga*.

Abstract Figure:
**Summary of *in situ* hybridization results:** genes coding for actors of NHEJ are expressed in the somatic nuclei and in the nurse nuclei of *Adineta vaga* individuals 2.5 hours post X-rays radiation, while genes coding for HR actors and PCNA (involved in multiple pathways including DNA replication and DNA repair: NER, BER, MR, HR) are expressed in the nurse nuclei 2.5 hours post radiation, and later in the maturing oocyte during oogenesis and in the laid eggs. Genes coding for actors highly expressed post-radiation, involved in the BER pathway appear to be only expressed in the somatic syncytium 2.5 hours post radiation, as well as the gene coding for the Ligase E, likely involved in DNA repair.

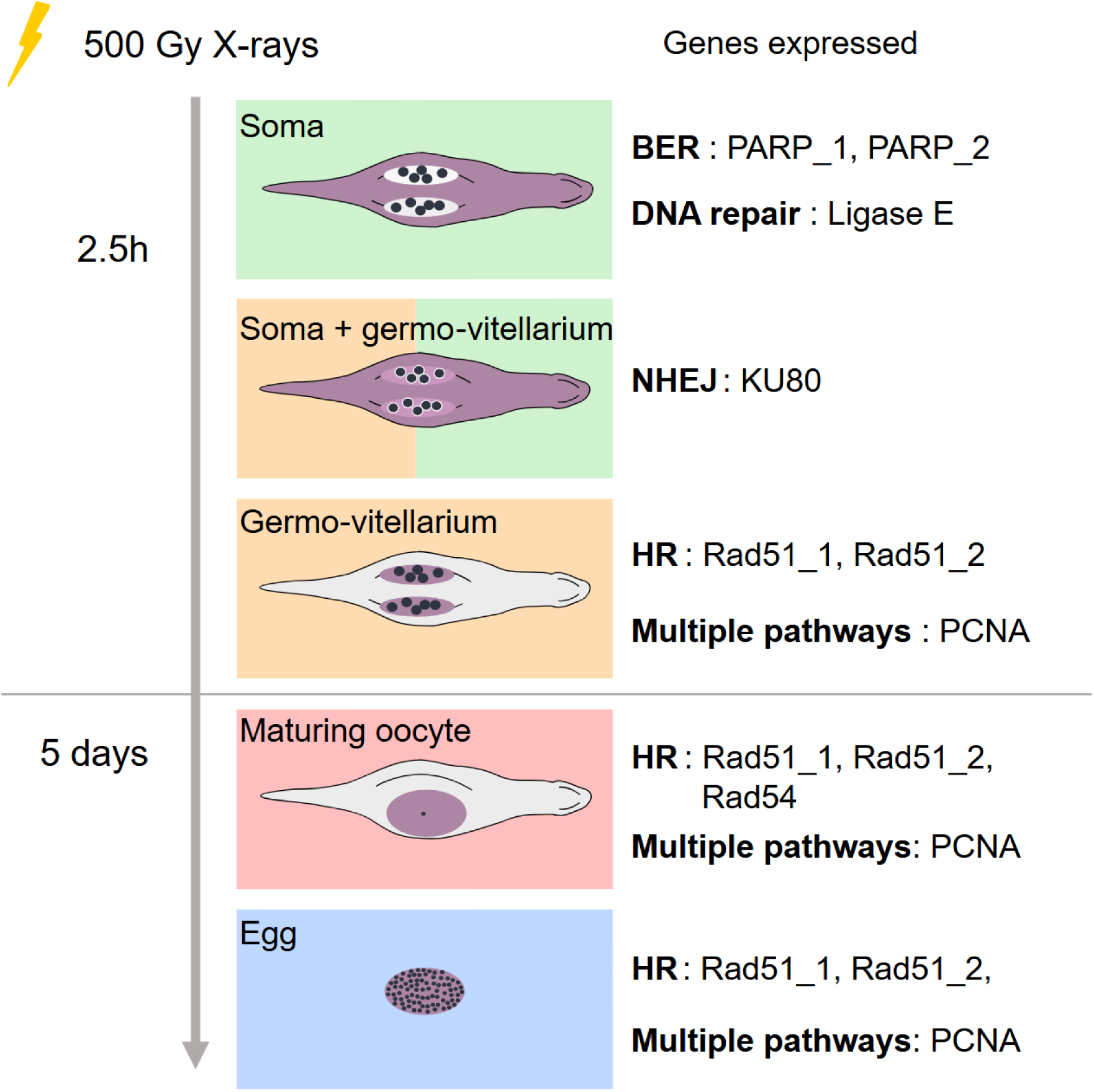

## Introduction

All organisms are susceptible to DNA damage caused by exogenous agents, including ultraviolet, ionizing radiation, environmental pollutants, and reactive oxygen species (ROS) generated by external oxidative stress, and by endogenous processes, such as errors introduced during DNA replication and ROS produced during aerobic respiration <u>(Cannan & Pederson, 2016; Jeggo & Löbrich, 2007)</u>. Exogenous agents like ionizing radiation are rare but harsh and can induce various types of DNA damage including double-strand breaks (DSBs), single-strand breaks (SSBs), mismatches, modified bases and abasic sites <u>(Zhao et al., 2020)</u>. DSBs are particularly critical for organismal survival due to their severe impact on genome integrity. If not properly repaired, DSBs can lead to chromosomal aberrations, translocations, or cell death, while mis-repair may result in mutations and genomic instability (<u>Jeggo & Löbrich, 2007; Chang et al., 2017; Zhao et al., 2020</u>).

Two main pathways are known to repair DSBs and appear conserved in eucaryotes: homologous recombination (HR) and non-homologous end-joining (NHEJ). While HR utilizes strand invasion and template-directed repair to produce a high-fidelity DNA repair, NHEJ consists of a re-adhesion of the two extremities of the DSB, being more error-prone <u>(Chatterjee & Walker, 2017)</u>. These two pathways rely on different molecular actors. Our understanding of these components is largely based on classical eukaryotic model systems, particularly yeast (*Saccharomyces cerevisiae*) and mammals. In eucaryotes, NHEJ involves the conserved KU70–KU80 heterodimer for DSB recognition, followed by the recruitment of the DNA-dependent protein kinase catalytic subunit (DNA-PKcs, kinase restricted to vertebrates) and end processing and ligation mediated by the XRCC4–DNA ligase IV complex <u>(Lieber, 2010; Chang et al., 2017; Zhao et al., 2020)</u>. HR relies on the conserved eukaryotic recombinase Rad51, assisted by mediator proteins (e.g., BRCA2, in vertebrates and plants) and accessory proteins such as Rad54, which promote homology search and strand invasion by promoting the interaction of Rad51-coated single-stranded DNA (ssDNA) with a homologous template, either the sister chromatid, or the homologous chromosome during meiosis <u>(von Nicolai et al., 2018, reviewed in Raina et al., 2025)</u>. Other types of DNA damages are being repaired by other pathways such as nucleotide damages by base excision repair (BER) (e.g., single base damages such as oxidation, deamination, and alkylation <u>(Chatterjee & Walker, 2017; Krokan & Bjørås, 2013; Beard et al., 2019)</u>, or nucleotide excision repair (NER) (e.g., bulky DNA lesions; <u>(Chatterjee & Walker, 2017; Schärer, 2013)</u>) which are broadly conserved across eukaryotes, although characterized mainly in the same model organisms. Although DNA repair pathways are specialized, the dynamic coordination of their factors often relies on scaffold proteins such as proliferating cell nuclear antigen (PCNA), which functions as a conserved eukaryotic sliding clamp and mediates the timely and precise recruitment of factors across multiple repair processes, including BER, NER, mismatch repair (MR), and HR (reviewed in Slade, 2018).

Organisms from a wide range of evolutionary clades, including some metazoan species, demonstrate notable or remarkable resistance to DNA damage caused by ionizing radiation. While mammalian cells can survive only relatively low doses of X-rays, typically with a lethal dose 50 (LD50) ranging from 2 to 6 Gy (Kohn and Kallman, 1957; Mole, 1984), radioresistant animals such as the microscopic bdelloid rotifer *Adineta vaga* can survive up to 5000 Gy of X-rays, although their reproductive rate decreases around 750 Gy <u>(Hespeels et al., 2020; Terwagne et al. 2022)</u>. Bdelloid rotifers are limno-terrestrial animals, are eutelic (i.e. fixed number of somatic cells when they reach maturity), and have a complete nervous, muscular, digestive and excretory system. Besides their extraordinary radiation resistance, they can survive complete desiccation and repeated freeze–thaw cycles. These environmental stresses induce, much like high doses of ionizing radiation, numerous DNA DSBs and an increased oxidative stress (Hespeels et al., 2014; Hespeels et al., 2023). Moreover, bdelloid rotifers represent a diverse animal clade reproducing exclusively parthenogenetically through a modified meiosis, not requiring any sperm or males for reproduction <u>(Ricci & Fontaneto, 2009; Segers, 2007; Welch et al., 2009, Terwagne et al. 2022)</u>.

Through pulse-field gel electrophoresis (PFGE) the progressive re-coalescence of sheared genomic DNA into high-molecular-weight molecules has been shown in the bdelloid rotifer *A. vaga* (<u>Hespeels et al., 2014, 2020</u>), confirming efficient somatic DNA repair within 24h following the damage (<u>Terwagne et al., 2022</u>). Time-resolved transcriptomic and proteomic surveys also revealed a rapid induction of the DNA-repair machinery: within 2.5 h after exposure/rehydration the transcripts of key base-excision repair (BER) factors (PARPs), canonical non-homologous end-joining (NHEJ) factors (KU80, Artemis, APLF) and DNA repair related actors (Ligase E) are highly upregulated (<u>Moris et al., 2024</u>), followed by an accumulation of their encoded proteins within 24 h (<u>Nicolas et al., 2023</u>). The PCNA gene, involved in multiple pathways (HR, BER and NER), is also strongly upregulated after radiation (<u>Moris et al., 2024</u>). This rapid molecular response correlates with the restoration of genome integrity (<u>Hespeels et al., 2014, 2020</u>).

While pulse-field gel electrophoresis (PFGE) reveals a DNA smear during the repair process, reflecting fragmented or partially repaired genomic DNA in somatic tissues (<u>Hespeels et al., 2014, 2020, 2023</u>), descendants of irradiated *A. vaga* no longer exhibit such smearing and display restriction patterns indistinguishable from the parental line (<u>Terwagne et al., 2022</u>). However, recent whole-genome sequencing analyses have uncovered a more complex and dynamic view of DNA repair in this species. Irradiated lineages exhibit long (up to several megabases) loss of heterozygosity (LOH) tracts flanked by heterozygous regions caused mostly by deleted fragments, together with progressive recovery of sequence coverage over successive generations. These patterns reflect a balance between DNA degradation and homologous recombination (HR)-mediated repair, generating subclonal heterogeneity in deletion size and repair extent (<u>Houtain et al., 2024</u>). Based on these observations, Houtain et al. (2024) proposed a transgenerational repair mechanism termed break-induced homologous extension repair (BIHER), in which fragmented chromosomes in *A. vaga* are transmitted through mitosis and progressively reconstructed via HR-mediated copying from the intact homolog, probably during the non-reductional meiosis when homologous chromosomes pair.

Taken together, available evidence suggests that *A. vaga* relies on rapid, KU-dependent non-homologous end joining (NHEJ), with auxiliary base excision repair (BER), in somatic cells, whereas DNA repair in the germline is delayed until oogenesis, when homologous chromosomes align and homologous recombination (HR) becomes feasible (<u>Terwagne et al., 2022; Houtain et al., 2024</u>). Bulk transcriptomic and proteomic datasets predominantly captured this somatic response, as germline nuclei represent only a minor fraction of cells in dense *A. vaga* cultures that must be pooled for such analyses (<u>Hespeels et al., 2014, 2020</u>). Consequently, the spatiotemporal dynamics of DNA repair in germline cells remain poorly characterized. Here, we test the hypothesis that DNA repair in irradiated *A. vaga* is spatially partitioned, with NHEJ and BER predominating in somatic nuclei, and HR in germline nuclei, and compare irradiated mothers and laid eggs. To address this, we developed an *in situ* hybridization (ISH) approach on cryosections using DIG-labelled probes and NBT–BCIP staining, enabling spatial and temporal analysis of DNA repair gene expression following irradiation. We focused on key components of the NHEJ pathway (KU80), the HR pathway (Rad51_1, Rad51_2, Rad54), and factors involved in multiple repair pathways like PCNA, but also Ligase E likely involved in *A. vaga* DNA repair (<u>Nicolas et al., 2023; Moris et al., 2024</u>).

## Results

### 1. Selection of reference genes and DNA repair actors

Using published RNAseq data <u>(Moris et al., 2024)</u>, we first examined which genes are highly expressed in hydrated and post-irradiation conditions in *A. vaga* (TPMs>500) to be used as reference genes to establish the protocol of *in situ* hybridization. We did choose three genes, coding for 60S ribosomal protein L5-like (FUN_002510-T1), Actin (FUN_025627-T1) and HSP82 (FUN_020850-T1) (***Fig. S1***), showing high expression levels in all conditions.

Second, we searched for DNA repair genes with high expression levels following radiation (TPMs>500), to test our protocol on genes expressed only post-irradiation. The aim is to identify their spatial expression within *A. vaga* adults, both in somatic and germline, including nurse cells, maturing oocytes (largest oocyte within the ovary with progressive growth and cytoplasmic accumulation), and laid eggs (mitotically developing embryos) from irradiated mothers (see ***abstract Figure***). We selected two genes reaching the highest expression levels post-radiation and with the highest log_2_foldchange <u>(Moris et al., 2024)</u>: one coding for PARP_1 (FUN_025694-T1) involved in the BER pathway, and one gene coding for the Ligase E (FUN_003353-T1) likely involved in DNA repair (***Fig. S1***). In addition, we also considered a gene coding for PCNA (FUN_027556-T1) being involved in BER, NER, HR, and in replication (Slade, 2018; ***Fig. S1***). We selected another gene with intermediate expression level post-irradiation involved in the BER pathway: PARP_2 (FUN_018903-T1; TPMs>200) (***Fig. S2***).

Third, we identified the expression levels of selected key genes involved in NHEJ (KU80: FUN_022394-T1) and HR pathways (Rad51_1: FUN_022601-T1, Rad51_2: FUN_006091-T1, Rad54: FUN_021207-T1). The selected gene coding for KU80 being overexpressed post-radiation <u>(Moris et al., 2024)</u> reached intermediate expression level (TPMs>100), similar to PARP_2 (***Fig. S2***). Genes encoding Rad51_1, Rad51_2, and Rad54 showed low expression levels (TPMs < 50) under all conditions (with or without radiation; <u>Moris et al., 2024</u>). Although Rad51_2 was overexpressed at 2.5 and 8 hours post-radiation, its overall expression level remained low (TPMs<50, ***Fig. S3***).

### 2. *In situ* hybridization of selected genes

#### 2.1. *In situ* hybridization of reference genes

The three genes used as reference genes, 60S ribosomal RNA, Actin, and HSP82 were found to be expressed around the nurse nuclei in the germo-vitellarium in hydrated *A. vaga* (***Fig. S4)***, while no signal was found, as expected, in the negative control using the sense probe (***Fig. S5***). Increased staining was observed in this region after radiation for the reference genes 60S and particularly for HSP82.

#### 2.2. *In situ* hybridization of Ligase E and BER associated actors

*In situ* hybridization of the gene coding for Ligase E, highly expressed in *A. vaga* after radiation and likely involved in DNA repair (<u>Nicolas et al., 2023; Moris et al., 2024</u>) and two genes coding for PARP (PARP_1; PARP_2) associated with the BER pathway, led to a strong signal in the somatic syncytial tissue of irradiated *A. vaga*, while the germo-vitellarium around the nurse nuclei remained unstained (***Fig. 1, Fig. S6***). No signal was observed in maturing oocytes during oogenesis in irradiated mother, nor in the laid eggs from irradiated mothers (***Fig. 1***). Unlike the reference genes, no signal of these genes was observed in the non-irradiated control (***Fig. 1***), in line with the calculated TPMs for these genes with values close to zero in non-irradiated *A. vaga* (***Fig. S1***).

**Fig. 1.**
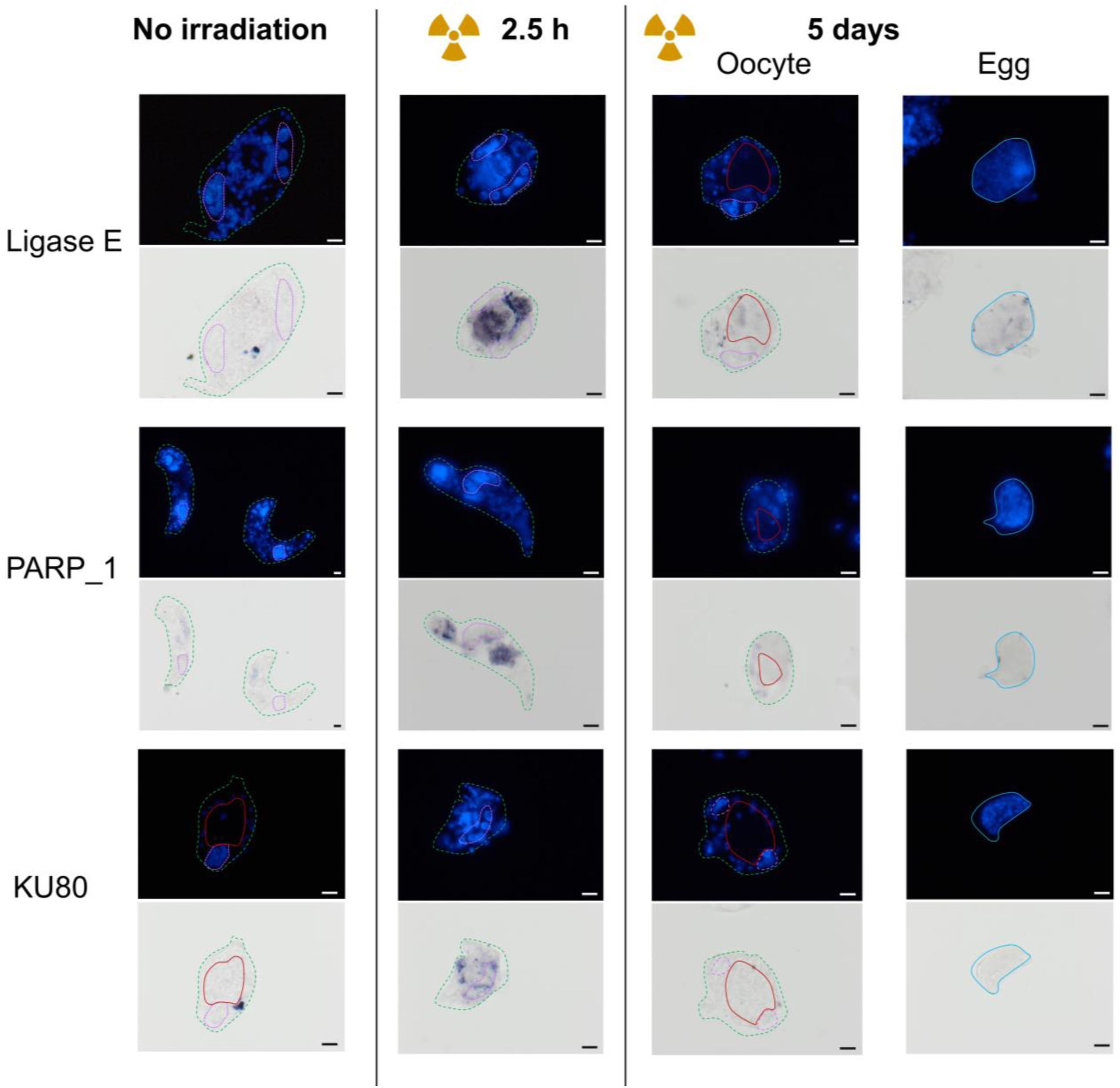
Spatial expression of Ligase E gene and genes involved in the BER (PARP_1) and NHEJ (KU80) repair pathways in cryosections of the bdelloid rotifer *Adineta vaga*. The first row represents the DAPI staining, the second row the probe staining through ISH. The dotted violet line surrounds nurse nuclei, while the continuous red line surrounds a maturing oocyte, the green dotted line surrounds the entire section of a rotifer individual, the blue line surrounds the laid egg. The first column represents the hydrated non-irradiated rotifer (control), the second column hydrated rotifer 2.5h post X-rays radiation (500 Gy), the third column hydrated rotifers 5-days post-irradiation with a developing oocyte, and in the fourth column a developing egg laid by a mother, 5 days post-irradiation.

#### 2.3. Genes involved in NHEJ

*In situ* hybridization experiments of the KU80 expressed gene gave a faint and dispersed staining in the somatic syncytial tissues in *A. vaga* rotifers irradiated with 500 Gy of X-ray (***Fig. 1***). Staining was also observed surrounding the nurse nuclei in the germo-vitellarium (***Fig. 1***, ***Fig. S6***). No staining was observed in rotifer oocytes’ undergoing oogenesis after radiation nor in the eggs laid by irradiated mothers (***Fig. 1***). Finally, no signal of KU80 was observed in the non-irradiated control (***Fig. 1***), in line with the calculated TPMs for that gene with values close to zero in non-irradiated *A. vaga* (***Fig. S2***).

#### 2.4. Genes involved in HR

The localization of the expression of three genes coding for proteins known to be involved in the HR pathway was investigated: Rad51_1, Rad51_2, and Rad54, as well as PCNA notably involved in HR, BER, NER, and DNA replication. The expressed genes, Rad51_1, Rad51_2, and PCNA, showed a similar spatial expression pattern: they were specifically detected in the germo-vitellarium of adults 2.5 hours post-radiation, 5 days post-radiation in maturing oocytes, and in eggs laid by irradiated mothers, while no signal was detected in the somatic syncytial tissue (***Fig. 2, Fig. S6***). Unlike the three genes above, the *in situ* hybridization targeting the investigated gene coding for Rad54 did not show any staining in the germo-vitellarium of adults 2.5 hours post-radiation or non-irradiated individuals nor in the somatic tissue. However, a faint signal of Rad54 is observed in maturing oocytes 5 days after irradiation and in eggs (***Fig. 2, Fig. S6***). The only gene observed to be expressed in non-irradiated *A. vaga* was Rad51_1, detected both in the maturing oocyte and in eggs laid by non-irradiated mothers (***Fig. 2, Fig. S6***). This is concordant with the calculated TPMs which show constant low expression of Rad51_1, also in non-radiation condition (***Fig. S3***).

**Fig. 2.**
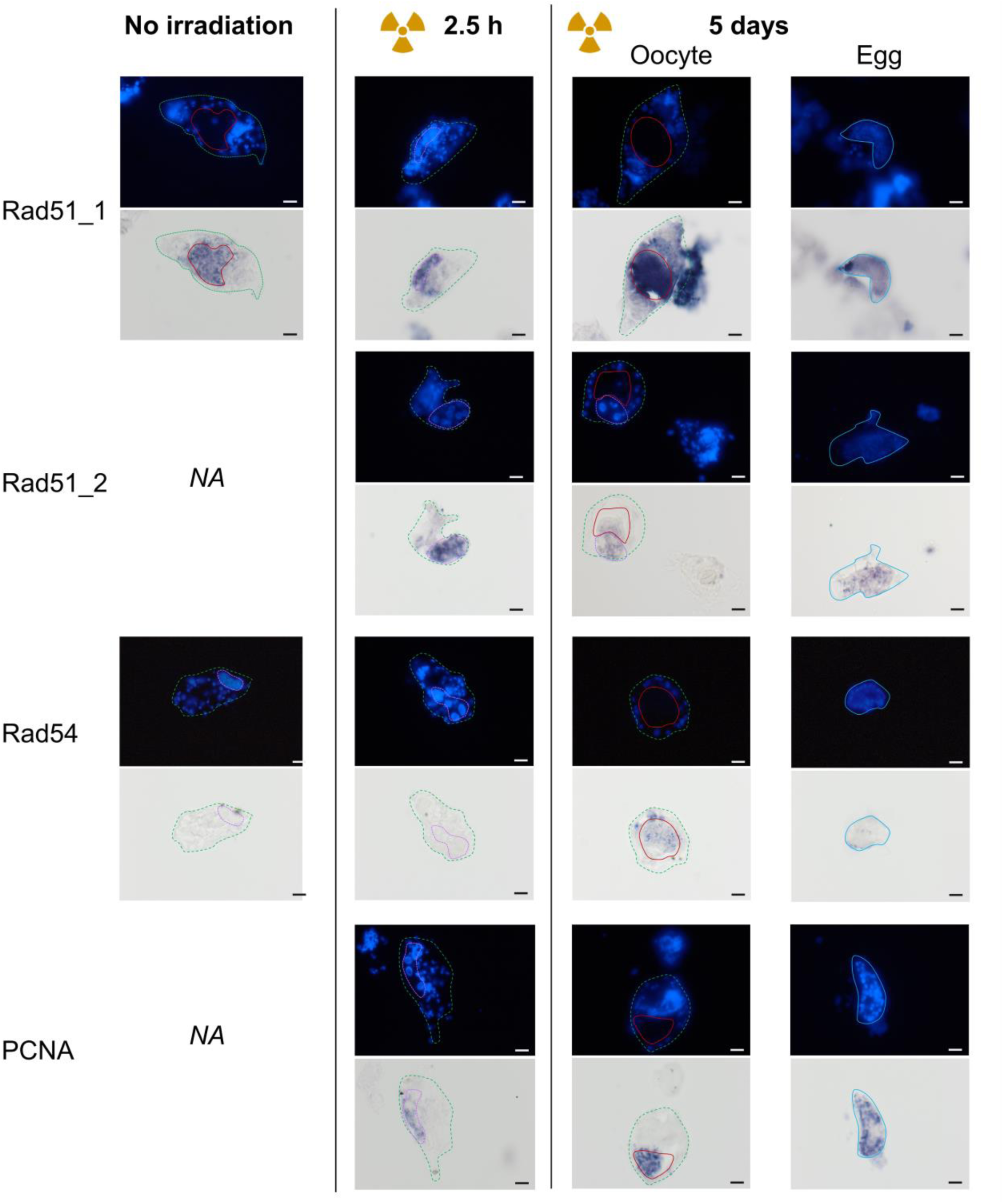
Spatial expression of genes involved in the HR repair pathway (Rad51_1, Rad51_2, Rad54) and PCNA in cryosections of the bdelloid rotifer *Adineta vaga*. The first row represents the DAPI staining, the second row the probe staining through ISH. The dotted violet line surrounds nurse nuclei, while the continuous red line surrounds a maturing oocyte, the green dotted line surrounds the entire section of a rotifer individual, the blue line surrounds the laid egg. The first column represents the hydrated non-irradiated rotifers (control), the second column hydrated rotifers 2.5h post X-rays radiation (500 Gy), the third column a hydrated rotifer 5-days post-irradiation with a developing oocyte, and in the fourth column a developing egg laid by a mother, 5 days post-irradiation.

## Discussion

Our newly established *in situ* hybridization method on cryosections of the bdelloid rotifer *A. vaga* provides a powerful tool to visualize the spatiotemporal expression of DNA repair pathway genes (and any other gene) separately in somatic and germline nuclei. By targeting mRNA, this approach enables the study of multiple genes at substantially lower cost than immunolabeling, which would require developing distinct antibodies for each protein of interest. Using this method, we provide the first spatial characterization of a few key DNA repair genes in *A. vaga* germ-line cells and maturing oocytes following irradiation, confirming distinct repair patterns between somatic and germline cells in bdelloid rotifer *A. vaga*.

### Somatic syncytium DNA repair via NHEJ and BER actors

Our study reveals the spatial expression of genes encoding key actors in Non-Homologous End Joining (NHEJ), such as KU80, Base Excision Repair (BER), including PARPs, and Ligase E likely involved in the BER pathway (Nicolas et al., 2023) within the somatic syncytium of *A. vaga* 2.5 hours post X-ray radiation exposure (***Fig 1***). These results align with previous transcriptomic and proteomic analyses of *A. vaga* (<u>Nicolas et al., 2023 and Moris et al., 2024</u>). By contrast, no probe signal was detectable in non-irradiated controls, indicating that basal expression is essentially silent and specifically induced by radiation, consistent with very low steady-state transcript levels under unchallenged conditions (<u>Moris et al., 2024</u>). No staining was observed for genes involved in the HR repair pathway in the somatic syncytium 2.5 hours post-radiation (***Fig 2***), reinforcing the hypothesis that BER and NHEJ are likely the predominant repair mechanisms in somatic tissues following radiation in *A. vaga*. The eutelic, post-mitotic nature of somatic syncytial nuclei in bdelloid rotifers favors the use of BER and NHEJ because these pathways operate efficiently without homologous templates <u>(Zhao et al., 2020)</u>. While NHEJ is potentially error-prone, its rapid action, together with BER, is sufficient to maintain somatic cell viability in *A. vaga* and support the synthesis of stress-response RNAs, repair proteins, antioxidants, and chaperones. Importantly, this occurs without compromising the genomes of future generations, as germline nuclei are spatially segregated and follow distinct repair dynamics (<u>Moris et al., 2024; Hespeels et al., 2020; Terwagne et al., 2022)</u>.

### Delayed HR repair in maturing oocytes from irradiated mother

Unlike somatic nuclei, oocytes of *A. vaga* undergo a modified meiosis, maturing one at a time within the mother <u>(Terwagne et al., 2022</u>; <u>Houtain et al., 2024)</u>. mRNA *in situ* hybridization shows robust expression of HR genes including Rad51_1, Rad51_2, Rad54, and PCNA, in maturing oocytes of irradiated mothers (***Fig. 2***), while NHEJ and BER genes remain undetectable (***Fig. 1***). Functionally, Rad51 promotes strand invasion and stabilizes replication forks, Rad54 facilitates chromatin remodeling and supports Rad51-mediated homologous recombination, and PCNA, a DNA sliding clamp, coordinates DNA synthesis and the recruitment of repair factors during homologous recombination and other DNA repair pathways <u>(Sebesta et al., 2013</u>; <u>Tavares et al., 2019)</u>. These results provide spatial evidence of the molecular machinery underlying HR in this system, transforming previously inferred genomic signatures into observable processes.

HR key actors have remained invisible in conventional transcriptomic and proteomic approaches, where signals from germline and somatic cells are pooled, masking germline-specific processes, particularly in maturing oocytes, which are generally underrepresented in standard cultures. Consequently, key homologous recombination (HR) actors remain largely undetected due to both cell-type dilution and the delayed activation of DNA repair during oocyte maturation. The protocol developed here provides two key improvements: (i) spatial resolution of DNA repair actor expression in maturing oocytes, and (ii) an increased proportion of maturing oocytes in boosted bdelloid rotifer cultures. The detection of these mRNAs in maturing oocytes 5 days post radiation suggest that DNA repair is delayed until HR occurs, enabling accurate repair of radiation-induced double-strand breaks during oogenesis, in agreement with F-ara-EdU labeling of prophase I oocytes (Fig. 2 <u>in (Terwagne et al., 2022))</u>. During modified meiosis I, HR is likely biased toward using the intact homologous chromosome as a repair template, as homologs are paired during prophase I (Fig. 3 in <u>(Terwagne et al., 2022))</u>, allowing strand invasion and crossover formation when required.

This germline-specific activation of HR actors provides a mechanistic basis for the patterns of loss of heterozygosity (LOH) and the linkage disequilibrium decay observed in bdelloid rotifers <u>(Simion et al., 2021; Vakhrusheva et al., 2020; Wilson et al., 2024; Houtain et al., 2024)</u>, and is consistent with the recently described Break-Induced Homologous Extension Repair (BIHER) pathway in *A. vaga* (<u>Houtain et al., 2024</u>). BIHER likely operates when a homologous template is accessible during modified meiosis I and relies on the HR-based machinery, including Rad51_1. HR in the germline plays a key role in maintaining chromosomal integrity, while BIHER acts across generations on larger deletions (<u>Houtain et al., 2024</u>), progressively enabling the restoration of a six homologous chromosome-pair karyotype. Our results here support a division of labor in DNA repair strategies in *A.vaga*, with rapid but potentially error-prone and faster NHEJ and BER pathways ensuring somatic survival, while delayed, high-fidelity HR in the germline preserves chromosomal integrity across generations.

### Dual DNA repair in nurse cells

In irradiated bdelloid rotifers, nurse cells express both HR (Rad51_a, Rad51_b) and NHEJ (KU80) genes as well as PCNA (***Figs. 1, 2***). These polyploid nuclei <u>(Amsellem & Ricci, 1982)</u> have been proposed to undergo endoreplication during oogenesis, as shown by strong F-ara-EdU DNA synthesis signals <u>(Terwagne et al., 2022)</u>, generating multiple identical genome copies while skipping canonical G1/G2/M phases. In this context, PCNA may support both DNA replication associated with endoreplication and repair-associated DNA synthesis, consistent with its roles in multiple DNA repair pathways, including HR. Polyploidization supports high biosynthetic activity, including yolk production and the synthesis of multiple mRNAs encoding factors required in the oocyte, which can be transferred to the oocyte via cytoplasmic bridges connecting the vitellarium syncytium to the germarium (<u>Amsellem & Ricci, 1982; Fontaneto & De Smet, 2014</u>).

HR and NHEJ gene expression in nurse nuclei may reflect a dual role: repair of irradiation-induced DNA breaks within nurse cells and potential maternal provisioning of transcripts and proteins to the oocyte. These may include HR-related factors required for DNA repair in the maturing oocyte, as well as proteins such as the heat shock protein HSP82, detected in these cells and potentially acting as a chaperone that stabilizes proteins or facilitates recruitment of HR factors <u>(Dubrez et al., 2020; Suhane et al., 2015)</u>. Co-expression of HR and NHEJ genes suggests that both pathways are available, with pathway choice likely influenced by DNA end resection status, cell cycle stage, and template availability. Extensive end resection promotes homologous recombination, whereas limited resection favors KU-mediated NHEJ (<u>Zhao et al., 2020</u>). The polyploid nature of nurse nuclei likely further shapes repair pathway usage by providing multiple homologous templates, thereby favoring HR. In mitotic cells, HR primarily uses sister chromatids during S/G2 phases, whereas inter-homolog recombination is constrained by the absence of pairing. Such high-fidelity repair may help limit deleterious mutations affecting essential functions, including DNA repair itself. At the same time, polyploidy increases tolerance to DNA damage through genomic redundancy (<u>Darmasaputra et al., 2024</u>). NHEJ may therefore act as a rapid backup pathway when templates are not immediately accessible, enabling nurse nuclei to maintain functionality and sustain oocyte provisioning, including transcripts for yolk proteins, HSP, meiosis, and DNA repair factors. Overall, nurse nuclei in *A. vaga* likely balance speed and fidelity by engaging NHEJ under urgent conditions while maintaining HR capacity to preserve genomic integrity under irradiation stress.

### HR repair in eggs from irradiated mothers

In eggs from irradiated mothers, the absence of detectable KU80, Ligase E, and PARP signals, together with the presence of HR-associated transcripts such as Rad51_1, Rad51_2, and PCNA (***Figs. 1, 2***), suggests that DNA repair preferentially relies on HR-related pathways rather than classical NHEJ. The detection of PCNA is consistent with its role in DNA replication in tissues undergoing active cell division, such as early embryonic stages, as well as in DNA repair-associated DNA synthesis. The repair bias towards HR pathway is consistent with the cellular context of early embryogenesis, in which accurate genome duplication is essential and homologous templates are readily available during replication. Early embryonic cells prioritize accurate genome duplication, and G1 phases are short or absent, leaving little opportunity for classical NHEJ, which is normally favored in G1. Even when DSBs are inherited from irradiated mothers, sister chromatids present during S and G2 provide physically proximal, perfectly homologous templates for repair, making homologous recombination (HR) or replication-coupled break-induced replication (BIR) the preferred pathways. One-ended breaks arising from stalled or collapsed replication forks, as well as locally damaged sister chromatids, are repaired via HR/BIR, and under extensive damage, alternative homologous sequences such as homologous chromosomes or repetitive/paralogous elements may also serve as templates. The coordinated action of Rad51, Rad54, and PCNA promotes strand invasion, chromatin remodeling, and repair DNA synthesis, enabling embryos to replicate through damaged DNA and maintain genomic integrity despite inherited lesions during early embryogenesis.

### Basal HR activity in non-irradiated oocytes and eggs

In non-irradiated conditions, Rad51_1 is observed in both developing maturing oocytes and eggs (***Figs. 2, S6***), consistent with a basal role in homologous recombination–associated genome maintenance during meiosis and DNA replication. In oocytes, Rad51 would mediate homologous recombination during meiotic DSB repair, where it assembles on resected single-stranded DNA and promotes strand invasion into a homologous template, typically the paired homologous chromosome. In addition to this canonical role, Rad51 likely also contributes to replication fork protection by stabilizing stalled replication intermediates under conditions of replication stress, limiting fork degradation and promoting restart when needed. In early embryos and eggs, Rad51 may similarly support genome integrity during rapid cell divisions by protecting and repairing DNA lesions arising during replication, including collapsed replication forks and associated repair intermediates. In mitotic cells, Rad51 has also been implicated in the protection of under-replicated DNA (<u>Wassing et al., 2021</u>), thereby contributing to faithful chromosome segregation and preventing genome instability in laid eggs.

Transcriptomic data indicate a functional divergence between Rad51 paralogs: one paralog, Rad51_1, is constitutively expressed under all conditions, likely performing routine replication-coupled repair, whereas Rad51_2 is induced only after irradiation of the mother (<u>Moris et al., 2024</u>; ***Fig. S3***), reflecting activation of the full HR repair program under stress. Other HR actors, such as Rad54 is largely absent under basal conditions (***Fig. 2***), either because their expression is very low or they are recruited only upon substantial DNA damage, explaining why signals are undetectable in normal oocytes and eggs.

## Conclusion

Our study supports the hypothesis that the mechanisms for repairing DSBs in *A. vaga* differ between germline and somatic lines. In the somatic syncytium, repair occurs rapidly through mechanisms like BER or NHEJ pathways, whereas DNA repair in germline nuclei is delayed until oogenesis, favoring template-dependent HR during oogenesis (***abstract Figure***). These observations extend our previous work on delayed germline repair in bdelloid rotifers (T<u>erwagne et al., 2022; Houtain et al., 2024</u>) by providing direct spatial evidence for differential deployment of DNA repair pathways between soma and germline. These observations can be placed in parallel with findings notably in *Caenorhabditis elegans*, where DNA damage in germline cells is repaired primarily through HR, whereas somatic cells can rely on both HR and NHEJ (<u>Lemmens & Tijsterman, 2011</u>). In that system, NHEJ appears to predominate in non-cycling somatic cells, while HR is more prevalent in proliferative cells, consistent with the cell-type-specific repair strategies observed here in *A. vaga*.

These observations also have important implications for long-term genome maintenance in bdelloid rotifers. HR-based repair generating LOH during meiosis in maturing oocytes may contribute to limiting mutation load, notably through the removal of deleterious alleles. Such repair may also promote allele conversion and structural genome changes, including duplications and deletions, in line with patterns previously described in irradiated lineages (<u>Houtain et al., 2024</u>). Although, in the absence of outcrossing, repeated LOH events may progressively increase homozygosity within lineages, HR repair in germline likely represents a key mechanism for repairing accidental DNA double-strand breaks encountered in the semi-terrestrial lifestyle of *A. vaga*, where desiccation can cause severe genome fragmentation (Hespeels et al., 2014). This strategy may therefore contribute to preserving genome integrity across generations exposed to recurrent desiccation–rehydration cycles in nature, while also providing a mechanistic framework for interpreting the response of *A. vaga* to irradiation-induced DNA damage in the laboratory.

## Methods

### *Adineta vaga* cultures

*Adineta vaga* cultures were maintained in Petri dishes (145 × 20 mm; Greiner Bio-One, Mosonmagyaróvár, Hungary) containing at least 15 mL of natural spring water (Spa®) and kept in thermostatic chambers at 21 °C. Cultures were fed three times per week with 500 µL of sterilized lettuce juice, and the water was renewed approximately once per week.

### *Adineta vaga* X-ray exposure

Approximately 100,000 rotifer individuals (corresponding to 4–6 dense cultures) were irradiated with 500 Gy of X-rays delivered at 8 Gy/min (225 kV, 19 mA) using a PXi X-Rad 225 XL irradiator (LARN, University of Namur, Belgium). A 2 mm aluminum filter was used to remove low-energy radiation. Rotifers were irradiated in Petri dishes (95 × 15 mm; Greiner Bio-One, Mosonmagyaróvár, Hungary) containing 15 mL of Spa® water. During irradiation, dishes were placed on a cooling block maintained at 4 °C to prevent heat-induced mortality.

### Boosting *Adineta vaga* in oogenesis

Approximately 100,000 *A. vaga* individuals were irradiated around midday. After irradiation, rotifers were fed with 200 µL of lettuce juice and maintained at 21 °C for the remainder of the day and the following two days. On day 4 post-irradiation, oogenesis was stimulated by increasing food availability (1000 µL lettuce juice) and temperature (25 °C). Between 24 and 36 h after this treatment, a higher proportion of the population displayed oogenesis, representing approximately one third of the individuals. At this stage, the population included *A. vaga* rotifers with or without developing oocytes, as well as laid eggs. Eggs were identified by the presence of numerous DAPI-stained nuclei and a visible outer layer corresponding to the egg envelope/cuticle in the brightfield channel (***Fig. S5***). Oogenesis was identified in DAPI-stained rotifers by the presence of a nucleus-poor cytoplasmic region within the body cavity, corresponding to the developing oocyte and its surrounding cytoplasm (***Fig. S5***).

### Studied genes

To establish the *in situ* hybridization protocol, we first selected three reference genes with high expression levels under both hydrated and post-irradiation conditions in *A. vaga* (reference genome used here is from <u>Simion et al., 2020</u>, and in <u>Moris et al., 2024</u>): 60S ribosomal protein L5-like (FUN_002510-T1), Actin (FUN_025627-T1), and HSP82 (FUN_020850-T1). After protocol optimization, we analyzed several DNA repair-related genes. These included Ligase E (FUN_003353-T1), PARP_1 (FUN_025694-T1), and PARP_2 (FUN_018903-T1), associated with the BER pathway, as well as PCNA (FUN_027556-T1), which is involved in DNA replication and multiple repair pathways, including BER, NER, and HR. We further examined KU80 (FUN_022394-T1), associated with NHEJ, and the HR-related genes Rad51_1 (FUN_022601-T1), Rad51_2 (FUN_006091-T1), two paralogs, as well as Rad54 (FUN_021207-T1).

### Probe synthesis

RNA probes were synthesized according to standard protocols <u>(Thisse & Thisse, 2008)</u> from cloned partial mRNA sequences of candidate genes using DIG RNA Labeling Kits (Roche, Basel, Switzerland). Strand cDNA was synthesized from isolated total RNA (1 µg, DNase-treated) of *A. vaga* for the genes 60S ribosomal protein, Actin, or from irradiated *A. vaga* 2.5h post 500 Gy X- ray radiation (when the expression was observed to be at its highest level: <u>(Moris et al., 2024)</u> for the genes Ligase E, Heat Shock Protein HSP82, KU80, Rad51_1, Rad51_2, Rad54, PARP_1, PARP_2 using Ambion RNAqueous®-4PCR Kit ( Fisher Scientific, Aalst, Belgium). Single strand cDNAs were synthesized with the SuperScript III First Strand Synthesis System (Fisher Scientific, Aalst, Belgium) and used to amplify target regions via polymerase chain reactions (PCRs) using the Phusion high fidelity kit (New England Biolabs, Ipswich, MA, USA). Amplicons were cloned with the NEB PCR Cloning kit pMiniT2.0 plasmid using the NEB® PCR cloning Kit (New England BioLabs, Massachusetts, USA). Plasmids were recovered through a miniprep using the GenEluteTM Plasmid Miniprep Kit (Sigma-Aldrich, Steinheim, Germany) or QIAprep® Spin Miniprep Kit (Qiagen, Hilden, Germany) and sent Genewiz (Genewiz, Leipzig, Germany) for sequencing. The nucleotide sequences of the plasmids were verified with Geneious <u>(Olsen et al., 2014)</u>. Linearization of the plasmids was obtained through separate digestion by BamH1-HF (New England BioLabs, Massachusetts, USA) or Not1-HF (New England BioLabs, Massachusetts, USA) enzymes, followed by a phenol-chloroform extraction.

The cloned gene fragments (92–825 bp long) were used to synthesize anti-sense and sense RNA probes using T7 (Roche, Mannheim, Germany) or SP6 (Roche, Mannheim, Germany) RNA polymerases. Probes were labeled with digoxigenin (DIG) using Digoxigenin-AP RNA labelling mix (Roche, Mannheim, Germany). A final clean-up of the probes was performed using RNeasy® MinElute® Cleanup Kit (Qiagen, Hilden, Germany). Sense probes served as negative control to each anti-sense probe. Oligonucleotide primers used for probe cloning (***Table 1***) were designed using Geneious <u>(Olsen et al., 2014)</u> and Primerblast <u>(Ye et al., 2012)</u>, and produced by IDT (Integrated DNA Technologies, Leuven, Belgium).

**Table 1.**
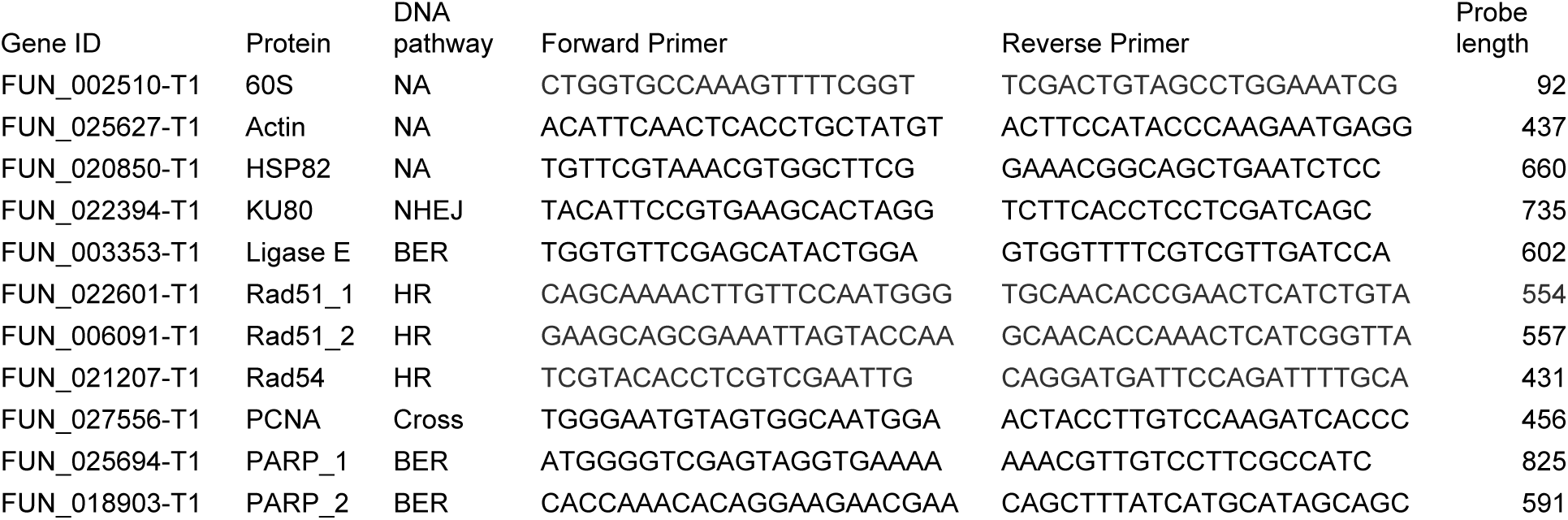
Studied genes, protein they are coding for and in which DNA repair pathway it is involved, nucleotide sequences of the designed forward and reverse primers used for the probe synthesis and the probe’s length.

Four to six dense cultures (i.e., 10 000 to 20 000 individuals per culture) of *A. vaga* were reassembled after centrifugations (10 minutes, 4000 rpm, 4 °C) to collect the rotifers as a pellet. This pellet was transferred into a 1,5 mL Eppendorf tube with the volume brought to 1 mL with Spa water (***Fig. S7***). The Eppendorf was centrifuged 1’30” at 6000 RPM at 4 °C and the supernatant was removed. The pellet (in the tube) was then put in liquid nitrogen (***Fig. S7***). Once removed, 1 mL of 4% paraformaldehyde (PFA) was added and samples were incubated for 1 hour at 4 °C to fix the rotifers (Fig. 9). PFA was removed by centrifugation and removal of the supernatant, then washed 2 times for 5 minutes in 1xPBS. Once washed and the phosphate buffered saline (PBS) removed, 1mL of 30% sucrose in 1xPBS was added to the pellet and incubated 1 hour at 4 °C (***Fig. S7***). Finally, sucrose was removed by centrifugation and the rotifer pellet was embedded using optimal cutting temperature (OCT) compound (Labonord S.A.S, Templemars, France) and stored at −80 °C (***Fig. S7***). Ten microns thick slices were then prepared the day of the hybridization using the Cryostat Leica CM1950 (Leica, Nussloch, Germany) (***Fig. S7***). Cryosections were fixed on the slides using 4% PFA and washed three times for 5 minutes using 1xPBS, then once with a solution of phosphate buffered saline and 0.1% Tween 20 (PTw) to remove all traces of PFA. Slides were dried, then washed three times for 5 minutes using PTw.

Depending on the investigated stages, we used hydrated rotifers, irradiated rotifers (2.5 hours post X-rays radiation), rotifers being boosted (5 days post radiation) or being boosted without radiation.

### *In situ* Hybridization

The *in situ* experiment is primarily based on the protocol made by Dearden et al. <u>(Dearden et al., 2009)</u>, with additional modifications from <u>(Moris et al., 2023)</u>. The slides were pre-hybridized by incubating them for 2h at 57 °C in a Dako Hybridizer (Agilent, Santa Clara, United States) with hybridization buffer (10% Dextran sulfate, 50% Formamide, 4X SSC, 1xDenhardt’s solution, 0.25 mg/ml tRNA, 0.5 mg/ml Heparin, 0.5% Tween 20). During the pre-hybridization, equal volumes of probes and carbonate buffer (120 mM Sodium Carbonate, 80 mM Sodium hydrogen carbonate, filled with Milli-Q water, pH 10.2) were mixed in a tube and probes were digested for 30 minutes at 60 °C. The slide’s hybridization buffer was replaced by a solution of 250 µL hybridization buffer and the digested probes (3 µL). Slides were incubated overnight at 57 °C using the HybEZ™ II Hybridization System (ACD - a Bio-Techne brand, Minneapolis, United States).

The next day, slides were washed 3 times for 30 minutes with the washing buffer (50% Formamide, 2X SSC, 0.5% Tween 20, filled with Milli-Q water) at 57 °C in a water bath and three times 5 minutes in PTw at room temperature. Blocking was performed by replacing PTw with a solution of PTw and 0.1% Bovine Serum Albumin (PBTw). After blocking with PBTw, the solution is replaced with a 1:1000 dilution of anti-DIG-AP antibody diluted in PBTw. The antibodies are then removed and replaced with a staining buffer (1/50 of NaCl 5M, 1/10 of Tris-HCl pH 9.5, 0.5% Tween 20, filled with Milli-Q water) which was incubated at room temperature in the dark because the reaction is light sensitive. Once staining is complete, tissues are washed twice for 5 minutes with PTw, then twice for 5 minutes with PBS to remove all traces of solutions and finally fixed with a solution of DAPI diluted 7500 times in Mowiol. Slides are stored at 4 °C in the dark.

The messenger RNA (mRNA) expression is detected using a chromogenic method, involving a combination of nitro blue tetrazolium (NBT) and 5-bromo-4-chloro-3-indolyl (BCIP), called NBT/BCIP (Roche, Mannheim, Germany). The signal was interpreted by comparing the staining obtained with antisense probes to that of the negative controls (sense probes; ***Fig. S6***).

### Imaging

Slides and pictures were taken using an automated fluorescence microscope BX63 (Olympus, Anvers, Belgium) with the Olympus cellSens Dimension software version 1.18 (Olympus, Anvers, Belgium). The blue precipitate resulting from the alkaline phosphatase reaction was observed using the brightfield channel of the microscope while the DAPI fluorescence was observed through the U-FUNA channel (excitation filter: 360-370, emission filter: 420-460). In both cases an Olympus sc50 color microscope camera was used.

## Author Contributions

Conceptualization VCM, BeH, KVD; Methodology VCM; Investigation VCM, AP, CH; Data Curation VCM, AP; Formal Analysis: VCM, AP; Writing – Original Draft VCM, AP, KVD; Writing – Review & edition VCM, AP, BH, BeH and KVD; Project Administration VCM, KVD; Funding Acquisition B.H. and KVD.

## Acknowledgements

The authors thank V Bielarz, J. Berthe, J. Lambert, M. Colinet, V. De Glas, A. Progneaux for their help with the establishment of the *in situ* hybridization protocol, and M. Terwagne and A. Houtain for sharing their knowledge and new results on *Adineta vaga* modified meiosis. They also thank A-C Heuskin for the use of the radiation infrastructure.

The authors thank the European Space Agency (ESA) and the Belgian Federal Science Policy Office (BELSPO) for their support in the framework of the PRODEX Program.

The research conducted here was also funded by a European Research Council (ERC) Consolidator Grant (RHEA) to K. Van Doninck.

## Supplementary Figures

**Fig. S1.**
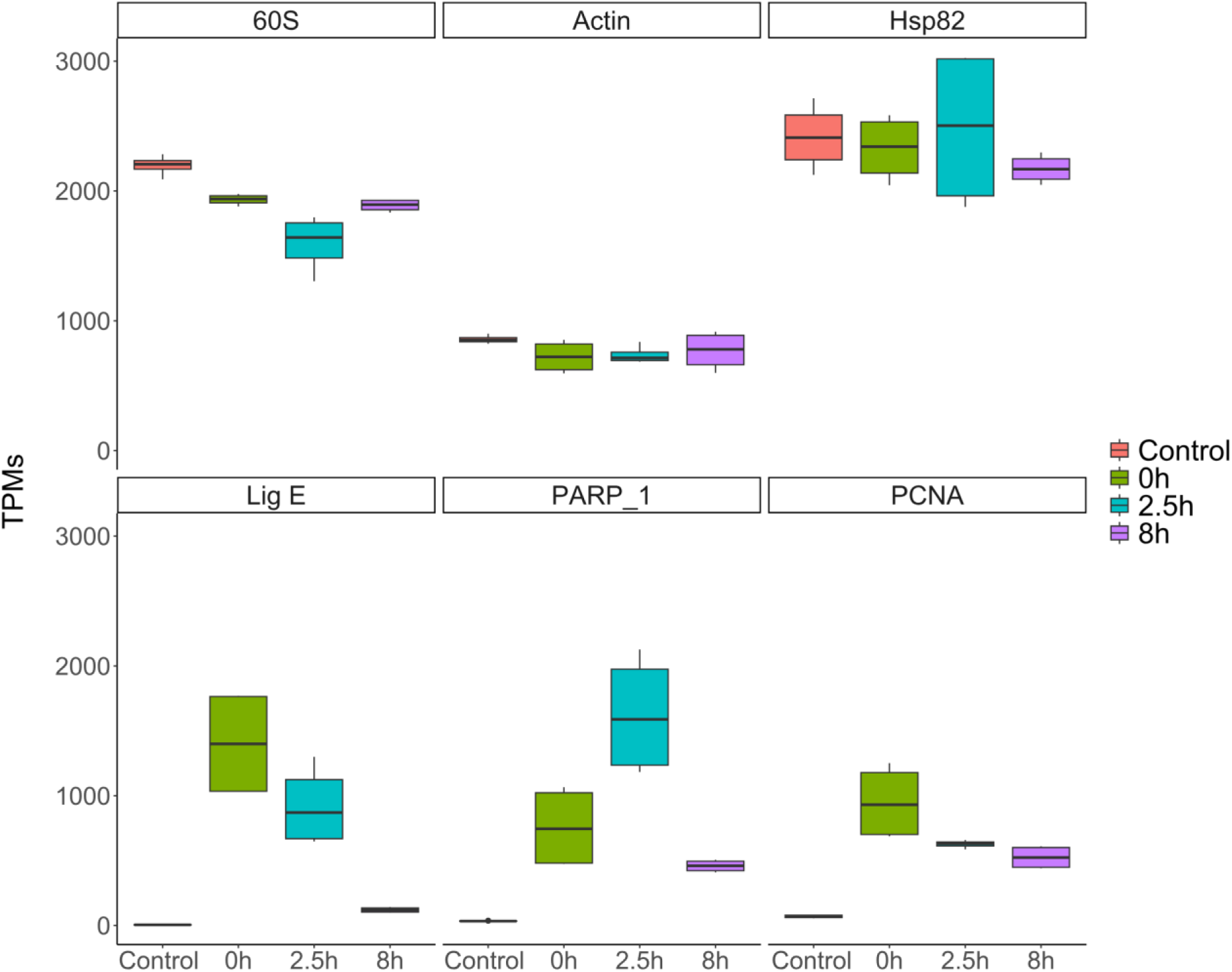
Boxplots of the expression levels (Transcript Per Millions: TPMs) of selected genes in bdelloid rotifer *Adineta vaga*. The top panel shows genes highly expressed (TPMs > 500) under hydrated control and irradiated conditions: 60S ribosomal protein L5-like (60S), Actin, and HSP82. The bottom panel displays genes that are highly expressed 0 and 2.5 hours post-X-ray radiation (500 Gy) but have close to null expression under hydrated control conditions: PARP_1 of the base excision repair (BER) pathway, Ligase E likely involved in DNA repair and PCNA, which is involved in multiple DNA repair pathways (BER, NER, and HR) and DNA replication. Transcriptomic data from Moris et al., 2024.

**Fig. S2.**
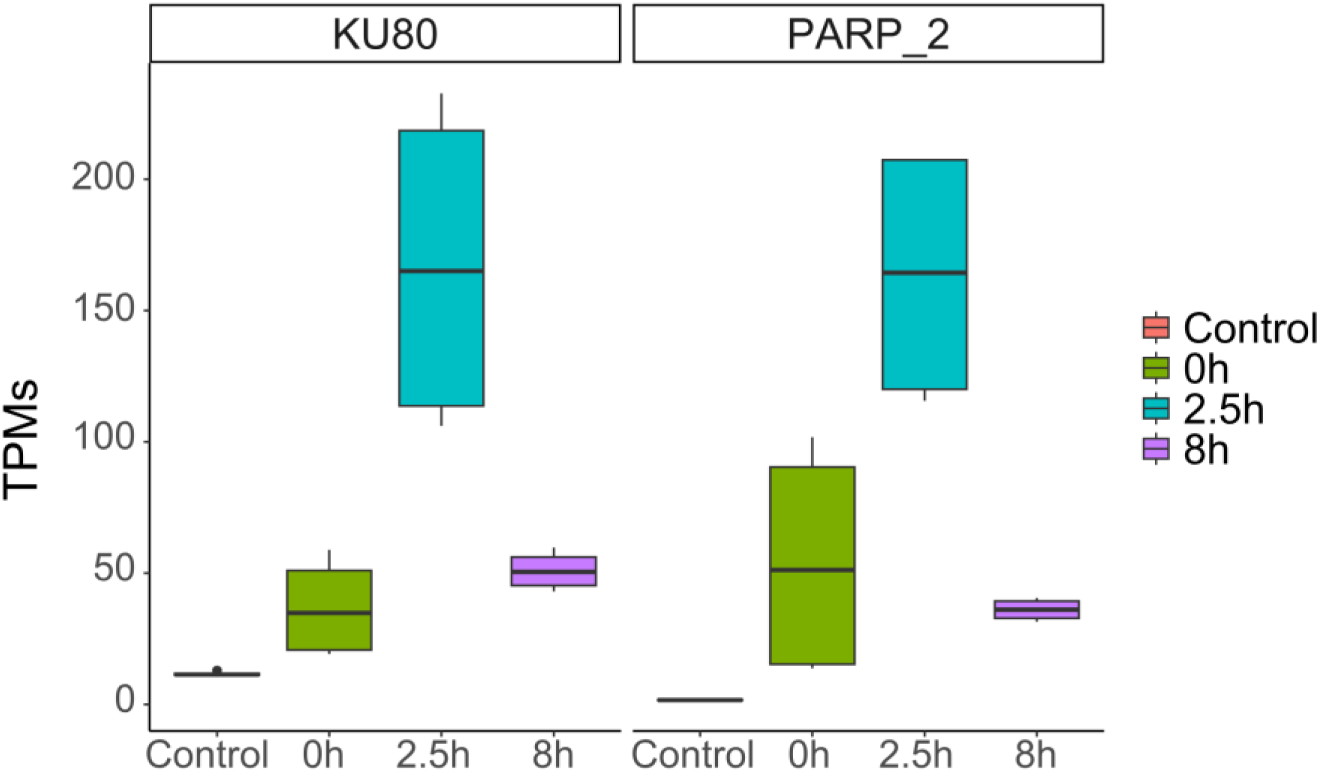
Boxplots of the expression levels (Transcript Per Millions: TPMs) of genes over-expressed and reaching high expression levels (TPMs>150) 2.5 hours post X-rays radiation (500 Gy): KU80 involved in NHEJ and PARP_2 involved in BER. Transcriptomic data from Moris et al., 2024.

**Fig. S3.**
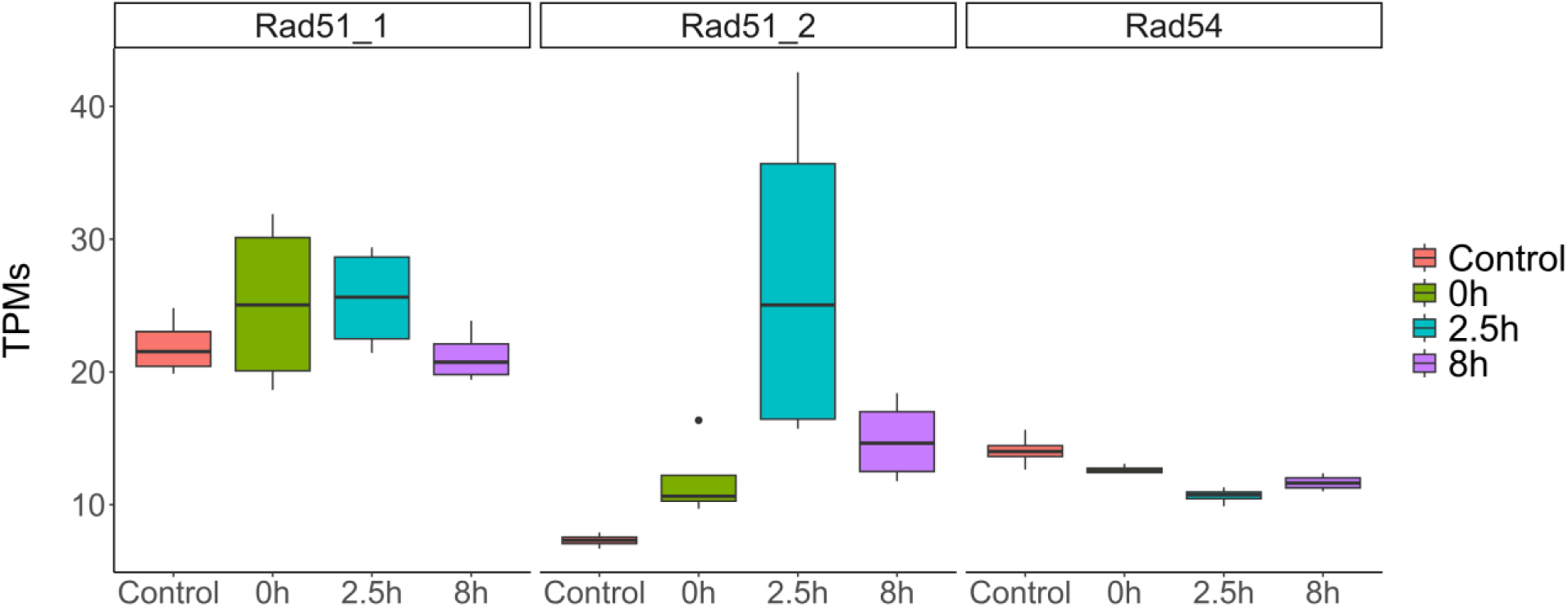
Boxplots of the expression levels (Transcript Per Millions: TPMs) of genes involved in HR: Rad51_1, Rad51_2, and Rad54 showing low expression levels (TPMs<50) in all conditions. Rad51_1 and Rad54 do not show any over-expression post X-rays radiation (500 Gy) whereas Rad51_2 shows a little over-expression 2.5 and 8 hours post radiation. Transcriptomic data from Moris et al., 2024.

**Fig. S4.**
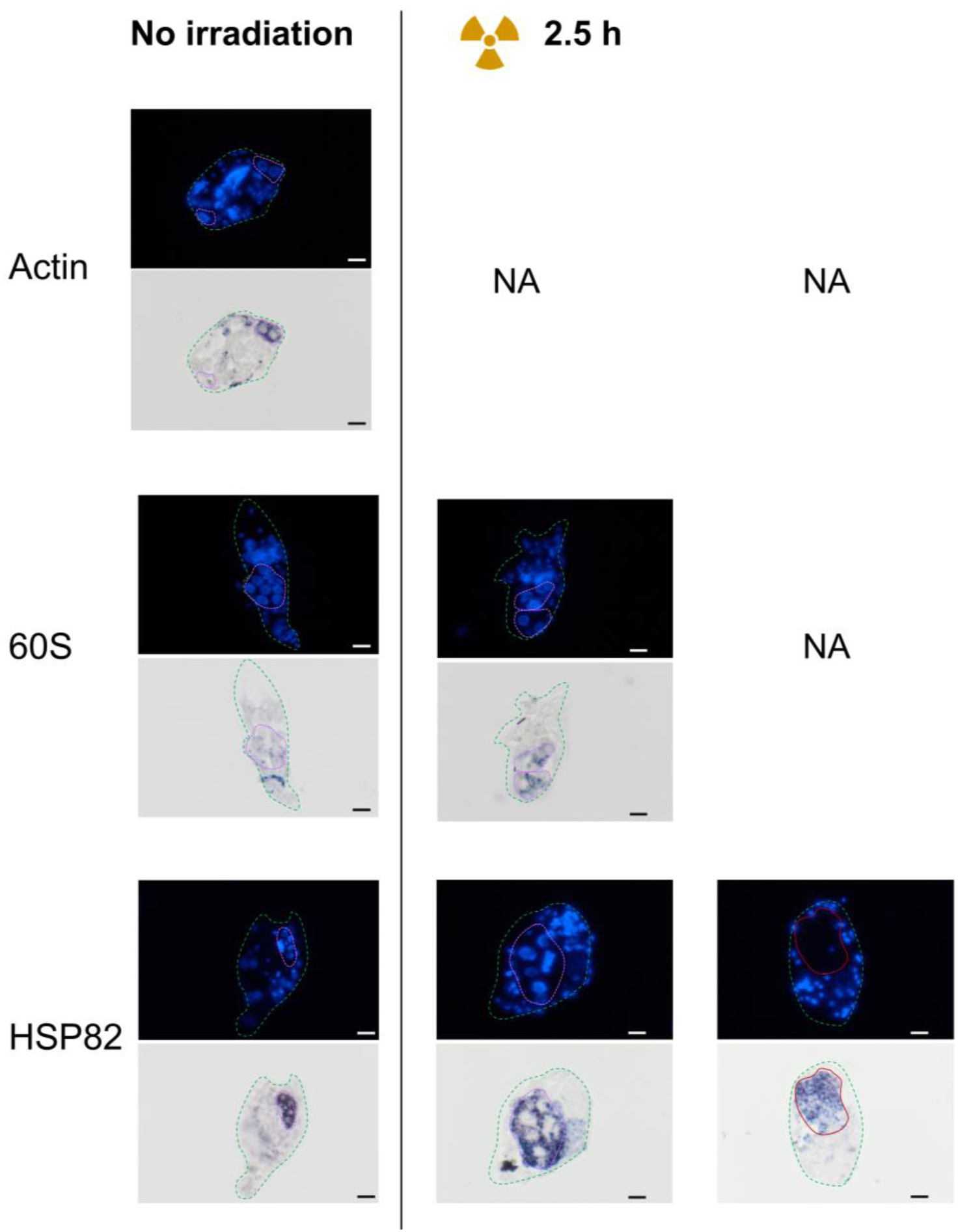
Spatial expression of reference genes coding for Actin, 60S ribosomal protein, and HSP82 in cryosections of bdelloid rotifer *Adineta vaga*. The first row represents the DAPI staining, the second row the probe staining through ISH. The dotted violet line surrounds nurse nuclei, while the continuous red line surrounds a maturing oocyte, the green dotted line surrounds the entire section of the rotifer individual. The first column represents the hydrated non-irradiated *A. vaga* adult rotifers and the second column hydrated *A. vaga* adult rotifers 2.5h post X-rays radiation (500 Gy). We highlighted in the third column rotifer individuals with a developing oocyte 2.5h post-radiation.

**Fig. S5.**
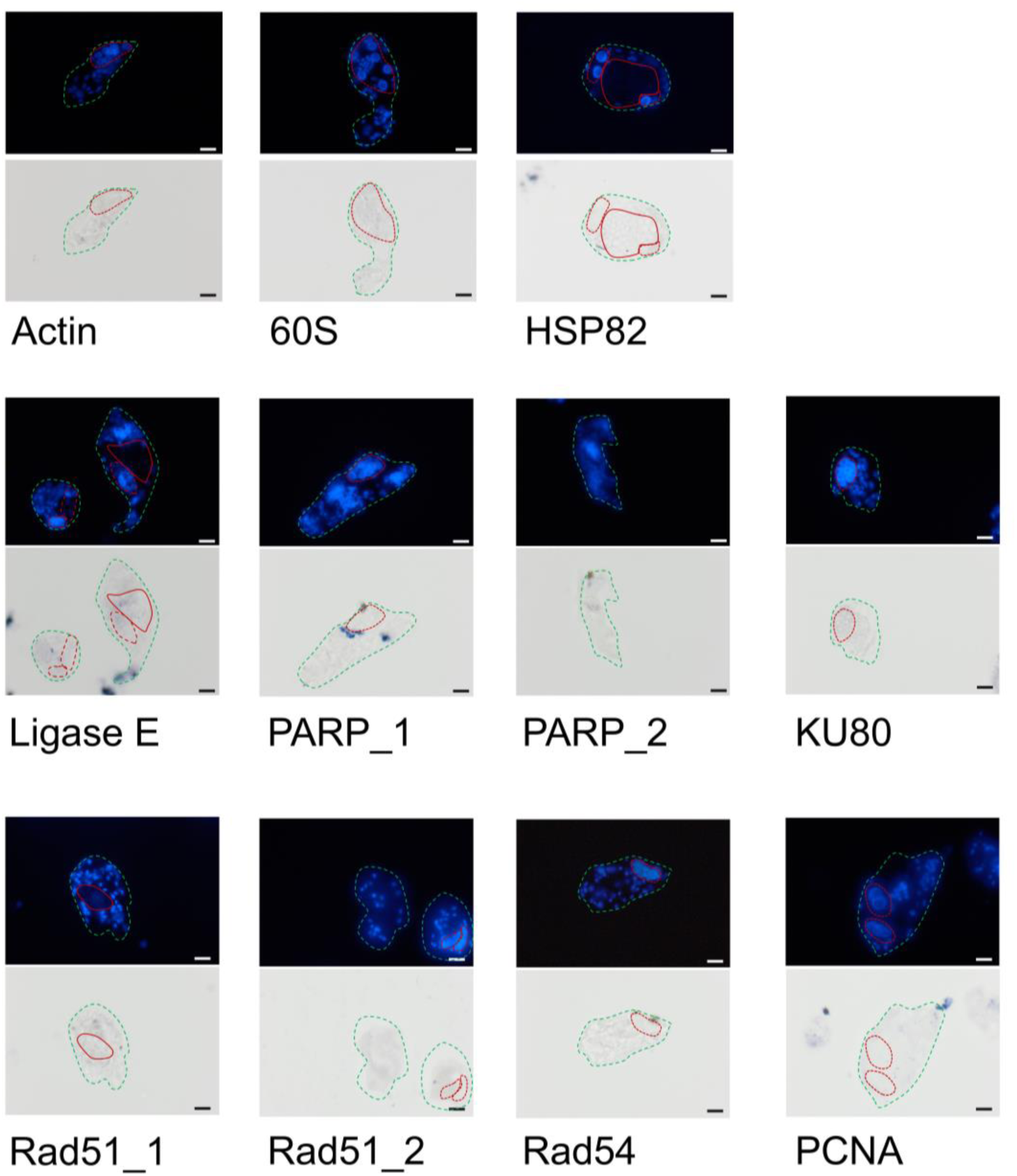
Negative control (sense probes, serving as negative control to each anti-sense probe) of the investigated genes on the bdelloid rotifer *Adineta vaga* (non-irradiated for the three reference genes, and 2.5h post radiation for DNA repair genes). The first row represents the DAPI staining, the second row the probe staining through ISH. The dotted violet line surrounds nurse cells, while the continuous red line surrounds a maturing oocyte, the green dotted line surrounds the entire section of the rotifer individual, the blue line surrounds a laid egg.

**Fig. S6.**
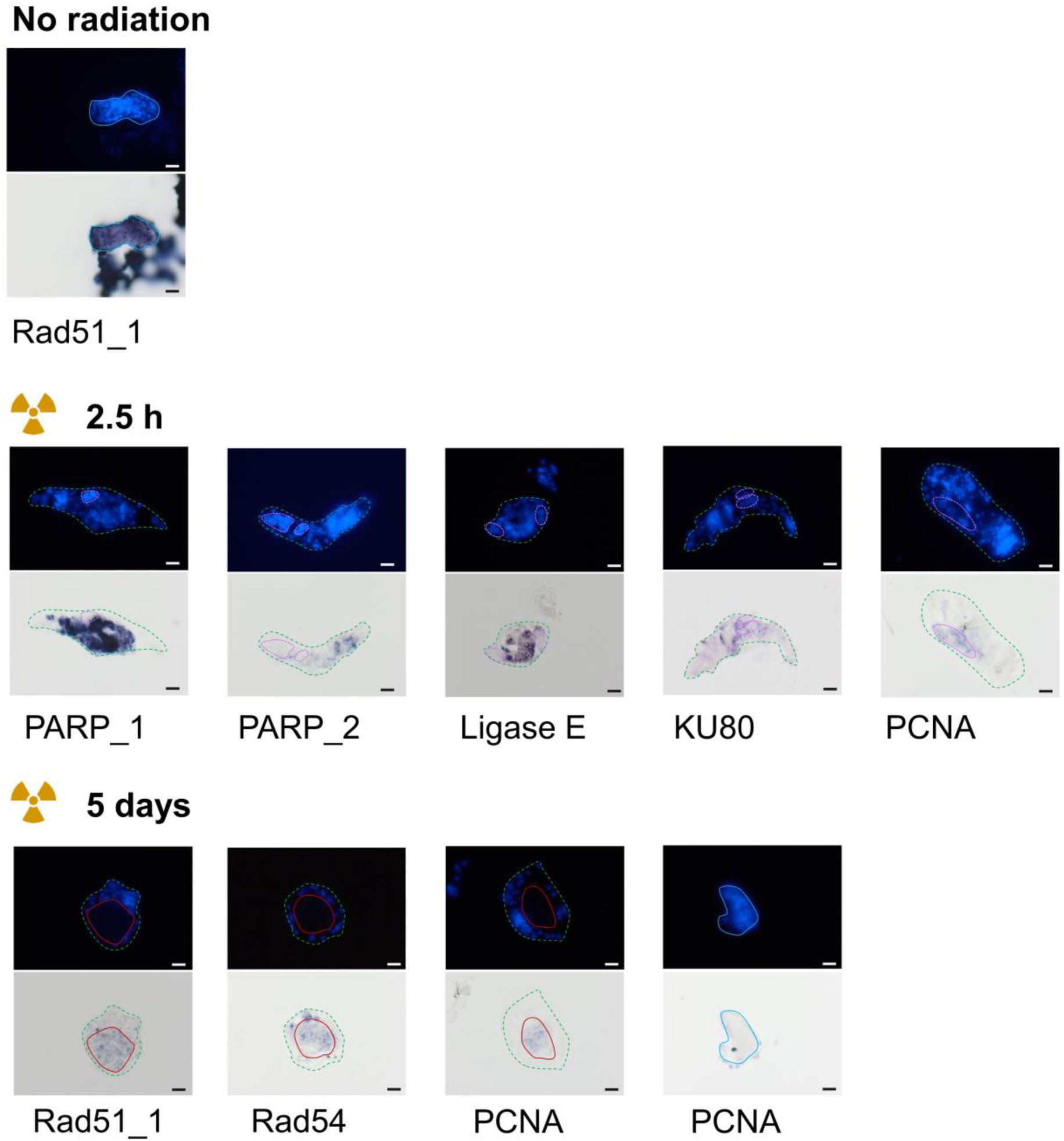
Additional results showing the spatial expression of investigated genes in cryosections of non-irradiated bdelloid rotifer *Adineta vaga*, in irradiated *A. vaga* 2.5 hours post radiation, and 5 days post radiation in *A. vaga* individuals presenting a maturing oocyte or in laid eggs (for PCNA gene). The first row represents the DAPI staining, the second row the probe staining through ISH. The first row represents the DAPI staining, the second row the probe staining through ISH. The dotted violet line surrounds nurse cells, while the continuous red line surrounds maturing oocyte, the green dotted line surrounds the rotifer, the blue line surrounds a laid egg.

**Fig. S7.**
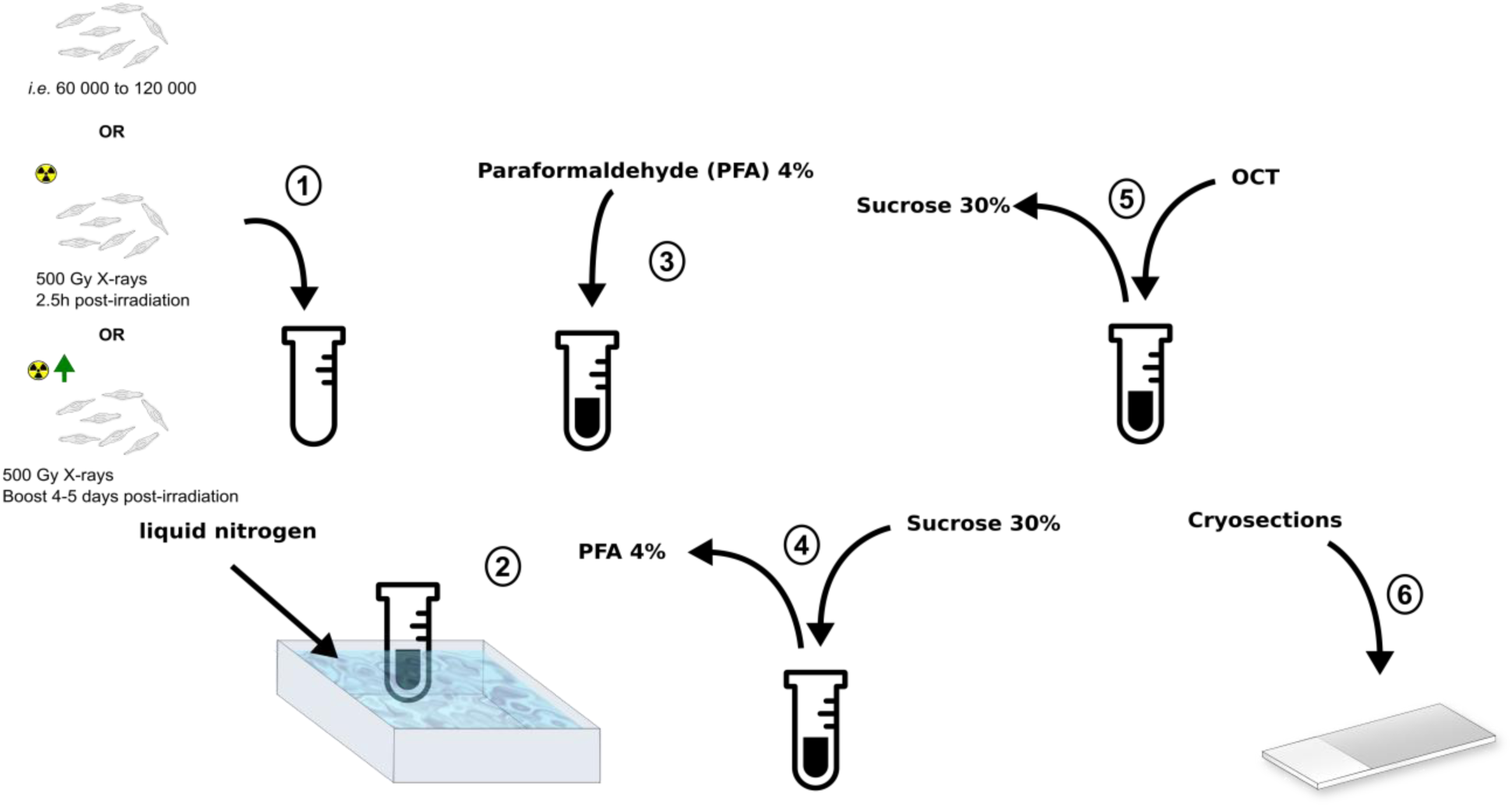
OCT embedding of *Adineta vaga*. (1) *A. vaga* pooling in an Eppendorf, (2) *A. vaga* pellet is instantly frozen in liquid nitrogen (3), the pellet fixed in paraformaldehyde (PFA) 4%, (4) PFA 4% replaced by sucrose 30%, (5) removal of sucrose and OCT added, (6) 10 µm cryosections on slides.

## References

Amsellem, J., & Ricci, C. (1982). Fine structure of the female genital apparatus of Philodina (Rotifera, Bdelloidea). Zoomorphology, 100(2), 89–105.

Beard, W. A., Horton, J. K., Prasad, R., Wilson, S. H. (2019). Eukaryotic base excision repair: New approaches shine light on mechanism. Annual review of biochemistry, 88:137–62

Cannan, W. J., & Pederson, D. S. (2016). Mechanisms and consequences of double-strand DNA break formation in chromatin. Journal of Cellular Physiology, 231(1), 3–14.

Chang, H. H. Y., Pannunzio, N. R., Adachi, N., & Lieber, M. R. (2017). Non-homologous DNA end joining and alternative pathways to double-strand break repair. Nature Reviews. Molecular Cell Biology, 18(8), 495–506.

Chatterjee, N., & Walker, G. C. (2017). Mechanisms of DNA damage, repair, and mutagenesis. Environmental and Molecular Mutagenesis, 58(5), 235–263.

Darmasaputra, G. S., van Rijnberk, L. M., & Galli, M. (2024). Functional consequences of somatic polyploidy in development. Development, 151(5), dev202392.

Dearden, P. K., Duncan, E. J., & Wilson, M. J. (2009). Whole-mount in situ hybridization of honeybee (Apis mellifera) tissues. Cold Spring Harbor Protocols, 2009(6), db.prot5225.

Dubrez, L., Causse, S., Borges Bonan, N., Dumétier, B., & Garrido, C. (2020). Heat-shock proteins: chaperoning DNA repair. Oncogene, 39(3), 516–529.

Fontaneto, D., & De Smet, W. H. (2014). 4. Rotifera. In Volume 3 Gastrotricha and Gnathifera (pp. 217–300). De Gruyter.

Hespeels, B., Knapen, M., Hanot-Mambres, D., Heuskin, A.-C., Pineux, F., Lucas, S., Koszul, R., & Van Doninck, K. (2014). Gateway to genetic exchange? DNA double-strand breaks in the bdelloid rotifer *Adineta vaga* submitted to desiccation. Journal of Evolutionary Biology, 27(7), 1334–1345.

Hespeels, B., Penninckx, S., Cornet, V., Bruneau, L., Bopp, C., Baumlé, V., Redivo, B., Heuskin, A.-C., Moeller, R., Fujimori, A., Lucas, S., & Van Doninck, K. (2020). Iron Ladies - How Desiccated Asexual Rotifer Adineta vaga Deal With X-Rays and Heavy Ions? Frontiers in Microbiology, 11, 1792.

Houtain, A., Derzelle, A., Llirós, M., Hespeels, B., Nicolas, É., Simion, P., Virgo, J., Heuskin, A.-C., Lenormand, T., Hallet, B., & Van Doninck, K. (2024). Transgenerational chromosome repair in the asexual bdelloid rotifer Adineta vaga. In bioRxiv (p. 2024.01.25.577190). 10.1101/2024.01.25.577190

Jeggo, P. A., & Löbrich, M. (2007). DNA dSsouble-strand breaks: their cellular and clinical impact? Oncogene, 26(56), 7717–7719.

Kim, J., Kim, J., & Park, C. G. (2016). X-ray radiation and developmental inhibition of Drosophila suzukii (Matsumura) (Diptera: Drosophilidae). International Journal of Radiation Biology, 92(12), 849–854.

Krokan, H. E., & Bjørås, M. (2013). Base Excision Repair. Cold Spring Harbor Perspectives in Biology, 5(4), a012583.

Lemmens, B. B. L. G., & Tijsterman, M. (2011). DNA double-strand break repair in *Caenorhabditis elegans*. Chromosoma, 120(1), 1–21.

Lieber, M. R. (2010). The mechanism of double-strand DNA break repair by the nonhomologous DNA end-joining pathway. Annual review of biochemistry, 79(1), 181–211.

Ma, C., Zhan, G., Zhong, Y., Liu, B., Gao, X., Xu, L., & Wang, Y. (2019). Effects of X-Ray Irradiation on the Eggs and Females of Dysmicoccus lepelleyi (Hemiptera: Pseudococcidae). Journal of Economic Entomology, 112(1), 134–138.

Moris, V. C., Bruneau, L., Berthe, J., Heuskin, A.-C., Penninckx, S., Ritter, S., Weber, U., Durante, M., Danchin, E. G. J., Hespeels, B., & Van Doninck, K. (2024). Ionizing radiation responses appear incidental to desiccation responses in the bdelloid rotifer Adineta vaga. BMC Biology, 22(1), 11.

Moris, V. C., Podsiadlowski, L., Martin, S., Oeyen, J. P., Donath, A., Petersen, M., Wilbrandt, J., Misof, B., Liedtke, D., Thamm, M., Scheiner, R., Schmitt, T., & Niehuis, O. (2023). Intrasexual cuticular hydrocarbon dimorphism in a wasp sheds light on hydrocarbon biosynthesis genes in Hymenoptera. Communications Biology, 6(1), 147.

Nicolas, E., Simion, P., Guérineau, M., Terwagne, M., Colinet, M., Virgo, J., Lingurski, M., Boutsen, A., Dieu, M., Hallet, B., & Van Doninck, K. (2023). Horizontal acquisition of a DNA ligase improves DNA damage tolerance in eukaryotes. Nature Communications, 14(1), 7638.

Olsen, C., Qaadri, K., Moir, R., Kearse, M., Buxton, S., Cheung, M., & Others. (2014). Geneious R7: a bioinformatics platform for biologists. International Plant and Animal Genome Conference Xxii. https://www.researchgate.net/profile/Richard-Moir/publication/268098577_Geneious_R7_A_Bioinformatics_Platform_for_Biologists/links/5717f17408ae30c3f9f17637/Geneious-R7-A-Bioinformatics-Platform-for-Biologists.pdf

Poinar, G. O., Jr, & Ricci, C. (1992). Bdelloid rotifers in Dominican amber: Evidence for parthenogenetic continuity. Experientia, 48(4), 408–410.

Ricci, C., & Fontaneto, D. (2009). The importance of being a bdelloid: Ecological and evolutionary consequences of dormancy. Italian Journal of Zoology, 76(3), 240–249.

Schärer, O. D. (2013). Nucleotide excision repair in eukaryotes. Cold Spring Harbor Perspectives in Biology, 5(10), a012609.

Sebesta, M., Burkovics, P., Juhasz, S., Zhang, S., Szabo, J. E., Lee, M. Y. W. T., Haracska, L., & Krejci, L. (2013). Role of PCNA and TLS polymerases in D-loop extension during homologous recombination in humans. DNA Repair, 12(9), 691–698.

Segers, H. (2007). Annotated checklist of the rotifers (Phylum Rotifera), with notes on nomenclature, taxonomy and distribution. Zootaxa, 1564(1), 1–104.

Simion, P., Narayan, J., Houtain, A., Derzelle, A., Baudry, L., Nicolas, E., et al. (2020). Homologous chromosomes in asexual rotifer *Adineta vaga* suggest automixis. bioRxiv, 2020-06.

Simion, P., Narayan, J., Houtain, A., Derzelle, A., Baudry, L., Nicolas, E., Arora, R., Cariou, M., Cruaud, C., Gaudray, F. R., Gilbert, C., Guiglielmoni, N., Hespeels, B., Kozlowski, D. K. L., Labadie, K., Limasset, A., Llirós, M., Marbouty, M., Terwagne, M., … Van Doninck, K. (2021). Chromosome-level genome assembly reveals homologous chromosomes and recombination in asexual rotifer *Adineta vaga*. Science Advances, 7(41), eabg4216.

Slade, D. (2018). Maneuvers on PCNA Rings during DNA Replication and Repair. Genes, 9, 416. 10.3390/genes9080416

Smith, J. M. (1986). Evolution: contemplating life without sex. Nature, 324(6095), 300–301.

Suhane, T., Laskar, S., Advani, S., Roy, N., Varunan, S., Bhattacharyya, D., Bhattacharyya, S., & Bhattacharyya, M. K. (2015). Both the charged linker region and ATPase domain of Hsp90 are essential for Rad51-dependent DNA repair. Eukaryotic Cell, 14(1), 64–77.

Tang, C. Q., Obertegger, U., Fontaneto, D., & Barraclough, T. G. (2014). Sexual species are separated by larger genetic gaps than asexual species in rotifers. Evolution; International Journal of Organic Evolution, 68(10), 2901–2916.

Tavares, E. M., Wright, W. D., Heyer, W.-D., Le Cam, E., & Dupaigne, P. (2019). In vitro role of Rad54 in Rad51-ssDNA filament-dependent homology search and synaptic complexes formation. Nature Communications, 10(1), 4058.

Terwagne, M., Nicolas, E., Hespeels, B., Herter, L., Virgo, J., Demazy, C., Heuskin, A.-C., Hallet, B., & Van Doninck, K. (2022). DNA repair during nonreductional meiosis in the asexual rotifer Adineta vaga. Science Advances, 8(48), eadc8829.

Thisse, C., & Thisse, B. (2008). High-resolution in situ hybridization to whole-mount zebrafish embryos. Nature Protocols, 3(1), 59–69.

Vakhrusheva, O. A., Mnatsakanova, E. A., Galimov, Y. R., Neretina, T. V., Gerasimov, E. S., Naumenko, S. A., Ozerova, S. G., Zalevsky, A. O., Yushenova, I. A., Rodriguez, F., Arkhipova, I. R., Penin, A. A., Logacheva, M. D., Bazykin, G. A., & Kondrashov, A. S. (2020). Genomic signatures of recombination in a natural population of the bdelloid rotifer Adineta vaga. Nature Communications, 11(1), 6421.

Raina, V. B., Jessop, A., & Greene, E. C. (2025). Biochemical mechanisms of genetic recombination and DNA repair. Annual review of biochemistry, 94 :161–193.

von Nicolai, C., Ehlén, Å., Martinez, J. S., & Carreira, A. (2018). Dissecting the Recombination Mediator Activity of BRCA2 Using Biochemical Methods. Methods in Enzymology, 600, 479–511.

Wassing, I. E., Graham, E., Saayman, X., Rampazzo, L., Ralf, C., Bassett, A., & Esashi, F. (2021). The RAD51 recombinase protects mitotic chromatin in human cells. Nature communications, 12(1), 5380.

Welch, D. B. M., Ricci, C., & Meselson, M. (2009). Bdelloid Rotifers: Progress in Understanding the Success of an Evolutionary Scandal. In I. Schön, K. Martens, & P. Dijk (Eds.), Lost Sex: The Evolutionary Biology of Parthenogenesis (pp. 259–279). Springer Netherlands.

Wilson, C. G., Pieszko, T., Nowell, R. W., & Barraclough, T. G. (2024). Recombination in bdelloid rotifer genomes: asexuality, transfer and stress. Trends in Genetics: TIG, 40(5), 422–436.

Ye, J., Coulouris, G., Zaretskaya, I., Cutcutache, I., Rozen, S., & Madden, T. L. (2012). Primer-BLAST: A tool to design target-specific primers for polymerase chain reaction. BMC Bioinformatics, 13, 134–134.

Zhao, L., Bao, C., Shang, Y., He, X., Ma, C., Lei, X., Mi, D., & Sun, Y. (2020). The Determinant of DNA Repair Pathway Choices in Ionising Radiation-Induced DNA Double-Strand Breaks. BioMed Research International, 2020, 4834965.

